# Multimodal optical imaging reveals spatial metabolic heterogeneity in the aging retina

**DOI:** 10.64898/2026.08.17.745175

**Authors:** Hongje Jang, Shuang Wu, Fangyuan Gao, Dorota Skowronska-Krawczyk, Lingyan Shi

**Author notes:** Correspondence to &.

## Abstract

Understanding how aging reshapes retinal metabolism requires methods that can resolve molecular and structural changes across the retina’s highly organized cellular layers. Here, we applied a nonlinear multimodal imaging platform that integrates fluorescence lifetime imaging microscopy (FLIM), second-harmonic generation (SHG), hyperspectral stimulated Raman scattering (HS-SRS), and deuterium oxide-based stimulated Raman scattering (DO-SRS) to map age-associated metabolic and compositional alterations in young and aged mouse retinas. FLIM analysis of the outer nuclear layer (ONL) revealed increased free NADH and NADPH fractions in aged retinas, consistent with reduced oxidative phosphorylation and enhanced lipid anabolic activity. SHG imaging of the sclera showed pronounced age-related remodeling of collagen organization, including increased fiber density, elevated anisotropy, and the emergence of densely crosslinked bundles in the central sclera. DO-SRS further demonstrated elevated lipid turnover in rod photoreceptor outer segments and the retinal pigment epithelium (RPE) with aging which was confirmed by lipidomic analysis. Complementary HS-SRS analysis revealed reduced triacylglycerol and cholesterol content together with localized sphingosine accumulation in the RPE. Together, these findings provide a spatially resolved view of metabolic remodeling in the aging retina and establish multimodal optical imaging as a powerful framework for studying alterations associated with age-related retinal disease.

---

Aging is accompanied by progressive physiological decline driven in part by metabolic dysfunction, which contributes to the onset and progression of many degenerative diseases(1, 2). Altered metabolism has been implicated in disorders affecting the nervous system, cardiovascular system, liver, and kidney, yet the molecular basis of these changes remains incompletely understood because of the complexity of tissue organization and cellular interactions(3–6). Defining how metabolism is spatially regulated during aging is therefore essential for understanding disease mechanisms and identifying therapeutic opportunities.

The retina provides a compelling system in which to address this challenge. As a highly specialized neural tissue, it operates under extreme metabolic demand and depends on intimate coupling between distinct cellular compartments, including photoreceptors, retinal pigment epithelial (RPE) cells, glia, and extracellular matrix-rich support structures(7–10). This layered organization sustains vision but also creates vulnerability: even subtle disruptions in local metabolism can propagate across compartments and compromise retinal function(10–12). Indeed, metabolic dysfunction has been implicated in major age-related ocular disorders, including glaucoma and age-related macular degeneration(13). However, despite growing recognition that retinal aging is fundamentally a metabolic process, its molecular reorganization across retinal space remains incompletely defined.

A major obstacle has been the difficulty of measuring metabolism within intact retinal architecture(14). Conventional biochemical assays average signals across heterogeneous cell populations and obscure layer-specific changes. Histological and molecular approaches have provided important insights into retinal degeneration(15), but they do not readily capture the dynamic chemical and metabolic states that precede overt structural damage. Recent spatial omics and advanced imaging methods have begun to reveal molecular organization in complex tissues, yet no single modality fully resolves the complementary dimensions of redox state, molecular composition, matrix organization, and biosynthetic activity that together define tissue metabolism(16, 17). Single imaging modalities capture only a limited range of metabolic activities; however, their spatial correlations can provide critical insight into the mechanisms of aging and disease. Because metabolic activities vary depending on subcellular location, precise localization of specific organelles is essential(18–20). For this reason, high-resolution optical multimodal imaging approaches offer a distinct advantage over techniques with lower spatial resolution, such as imaging mass spectrometry(21).

Multimodal optical imaging offers a route to bridge this gap. Fluorescence lifetime imaging microscopy (FLIM) can report intracellular redox state through endogenous metabolic cofactors(22–24). Second-harmonic generation (SHG) enables label-free visualization of non-centrosymmetric structures such as fibrillar collagen(25–28). Hyperspectral stimulated Raman scattering (HS-SRS) provides chemically specific maps of lipids and other biomolecules, whereas deuterium oxide-based stimulated Raman scattering (DO-SRS) measures newly synthesized macromolecules in situ(29–32). Integrated within a single framework, these complementary modalities make it possible to interrogate metabolism across multiple biochemical and structural dimensions while preserving spatial context(19, 33–35).

To determine how aging reshapes the retinal metabolic landscape, we performed multimodal optical imaging of young and aged mouse retinas using an integrated platform combining FLIM, SHG, HS-SRS, and DO-SRS. This approach captures multiple orthogonal features of tissue physiology, including redox state, collagen organization, chemical composition, and biosynthetic activity, within intact retinal architecture. Rather than a uniform decline, aging produced distinct and spatially restricted changes across retinal layers and adjacent support tissues, including altered redox balance in photoreceptor nuclei, remodeling of scleral and choroidal collagen, and layer-specific shifts in lipid composition and turnover in the outer retina and RPE. These findings identify retinal aging as a spatially organized process of metabolic remodeling and provide a framework for resolving localized vulnerabilities in age-associated ocular degeneration.

## RESULTS

### Multimodal imaging of retinal samples at two different ages

All experiments were performed on albino animals housed under a standard 12-hour light (<150 lux)/12-hour dark cycle. Images of mouse retinal tissue sections were acquired using a nonlinear multimodal imaging approach. As illustrated in Fig. 1a, second-harmonic generation (SHG), multiphoton fluorescence (MPF), and hyperspectral stimulated Raman scattering (HS-SRS) images were sequentially captured from each region of interest (ROI). In addition, newly synthesized lipids and proteins were isotopically labeled with deuterium by supplying heavy water (D_2_O) to the mice, enabling quantitative comparison of the ratio of newly synthesized to pre-existing molecules via SRS signal measurements. Following the order described in Fig. S1, from the multimodal image stack, various metabolic signals were analyzed.

**Fig 1.**
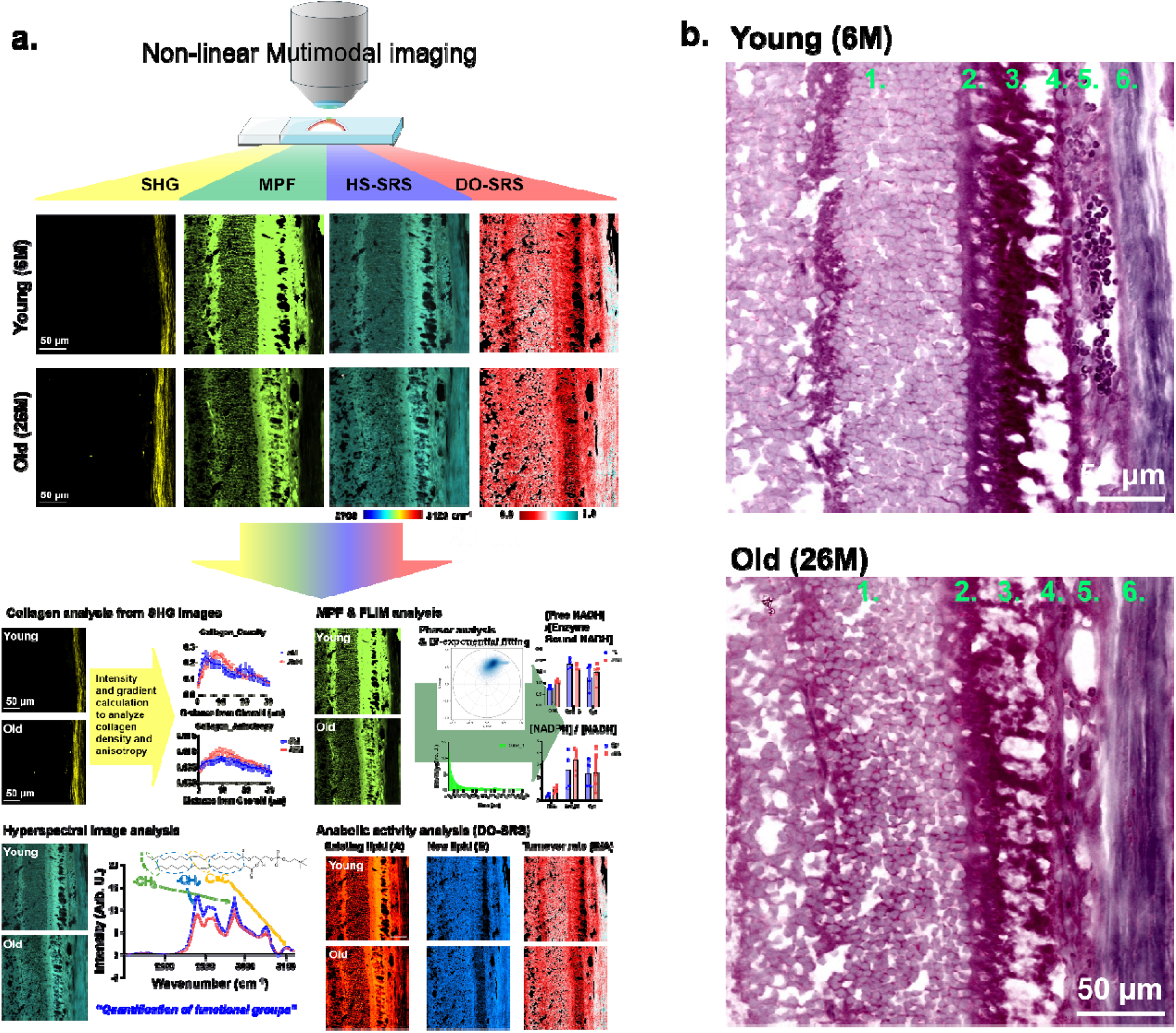
Schematic of collective chemical and metabolic study in retina. **a.** Retina samples of two different ages were studied using non-linear multimodal imaging setup of SHG, MPF, hyperspectral SRS, and DO-SRS. From different imaging channels, aging related molecular compositional, structural, and metabolic changes were measured. **b.** To segment tissue compartments, pseudo H&E images were generated from the hyperspectral images. (1. Outer Nuclear Layer, 2. Rod cell – inner segment, 3. Rod cell – Outer segment, 4, RPE layer, 5. Choroid, 6. Sclera)

The retina is a highly stratified structure comprising multiple distinct cellular layers. As shown in Fig. 1b, pseudo-hematoxylin and eosin (H&E) imaging was used to delineate these specific tissue compartments. Optical signals were selectively acquired and averaged from the outer nuclear layer (ONL), the inner and outer segments of the rod photoreceptor cells, the retinal pigment epithelium (RPE) layer, the choroid, and the sclera, in order to evaluate localized metabolic and chemical variations. The metabolic differences detected through these quantitative measurements are summarized in Fig. 1c.

### Attenuated energy production via oxidative phosphorylation in photoreceptors

Fluorescence lifetime imaging microscopy (FLIM) is a well-established technique for probing the local chemical environment of fluorophores. Even for chemically identical species, fluorescence lifetimes can vary substantially depending on the surrounding molecular environment. In retinal tissue, measurement of the autofluorescence lifetime of NADH provides critical insights into the relative activity of specific NADH-dependent metabolic pathways. Notably, free NADH, enzyme-bound NADH, and NADPH each exhibit distinct fluorescence lifetimes(22–24). As detailed in Fig. S2, these lifetime components can therefore be used to determine the relative proportions of free NADH, enzyme-bound NADH, and NADPH.

Because the outer segments of rod photoreceptor cells and erythrocytes within the choroid generate strong autofluorescence signals, these regions were excluded from the lifetime analysis. Among the remaining tissue compartments, the ONL exhibited the most pronounced age-related changes in the ratios of free NADH, enzyme-bound NADH, and NADPH. Specifically, the ONL of retinal tissues from aged mice demonstrated elevated levels of both free NADH and NADPH relative to those from young mice (Fig. 2a-d). In the inner segments of rod cells from aged mice, a trend toward decreased free NADH and increased NADPH was also observed, although it did not reach statistical significance. These findings are indicative of attenuated energy production via oxidative phosphorylation (OXPHOS)(36), accompanied by increased lipid anabolic activity(37).

**Fig 2.**
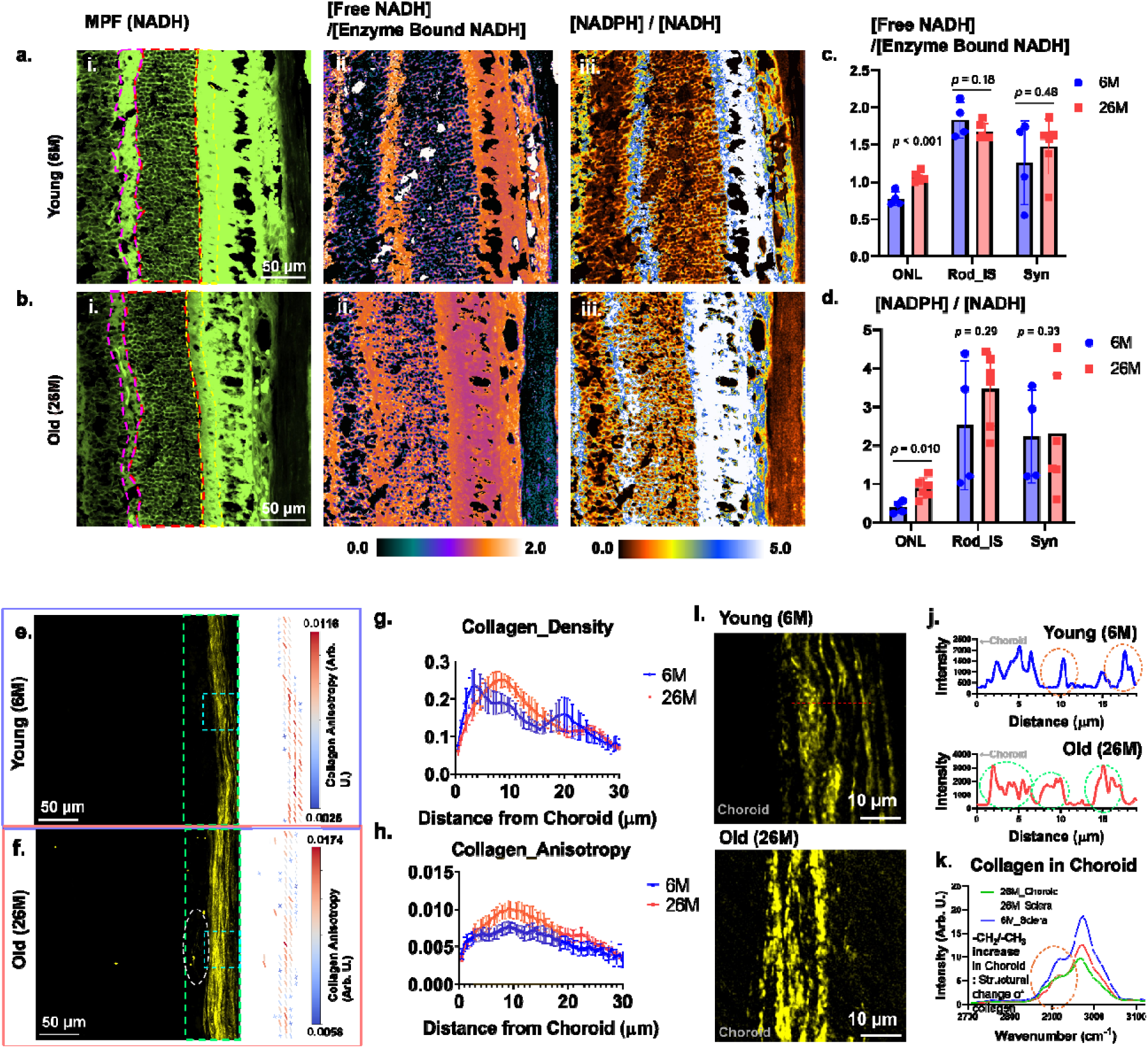
**a and b.** Fluorescence lifetime signals from two different aged samples were analyzed. i) Fluorescence intensity images of NADH channel. Synaptic layers (magenta dotted lines), outer nuclear layers (red dotted lines), and rod cell inner segments (yellow dotted lines) were analyzed. ii) Young sample shows relatively low amount of Free NADH level than old sample in nuclear layer. iii) Young sample has lower amount of NADPH than the retina samples from old mice. **c.** Averaged intensity values of Free NADH level was significantly increased in ONL. In inner segments of rod cells, the change was not significant enough, but the change had a tendency toward reduction. **d.** Averaged intensity values of NADPH level was significantly increased in ONL. **e and f.** Using SHG signal, collagen fiber distribution and structure in sclera were analyzed. Vectors of collagen fibers (in green dotted rectangles) were plotted with anisotropy information. **g and h.** Distance dependencies of Density and anisotropy from choroid are plotted. Averaged densities in two different sample groups were similar. Old retina has clear range with a high collagen density, around 10 microns. In the region of the high collagen density, old retina has relatively higher level of collagen anisotropy. **i and j.** Super-resolution SHG images from cyan dotted rectangles of panel a and b show the structural change of collagen fibers. In young retina sample, thin layers (white dotted ellipsoids) were detected, and the layers were parallelly aligned. In old retina sample, intensity and thickness increase (green dotted ellipsoids) were detected. **k.** Raman spectrum from sclera and choroid of aged mice exhibited different structural information. Ratio of –CH_2_/–CH_3_ Raman signals from young mice was higher than the signal ratio from aged mice. This ratio change implies collagen structural difference between two ROIs.

### Age-related remodeling of scleral and choroidal collagen structure

The sclera provides essential mechanical support to the eye, a property primarily governed by its dense collagen fiber network. To investigate age-related changes in scleral collagen fiber remodeling, SHG imaging was performed on tissue sections from two age groups. The acquired SHG images (Fig. 2e and f) were analyzed to quantify collagen fiber density and anisotropy. Following the methodology described in Fig. S3, the spatial distributions of collagen density and anisotropy were plotted as a function of distance from the choroid (Fig. 2g and h).

In tissues from young mice, the region of highest collagen fiber density was located in close proximity to the choroid (approximately 3 µm), whereas in aged mice, this peak density region shifted away from the choroid into the deeper scleral layers. Furthermore, tissues from young animals exhibited a substantially more uniform anisotropy distribution, while those from aged animals displayed an anisotropy profile that closely mirrored the density distribution. Given that anisotropy reflects structural heterogeneity, these results indicate the formation of highly dense, strongly crosslinked collagen fiber bundles within the middle layer of the aging sclera. Super-resolution SHG imaging further corroborates these age-related structural alterations (Fig. 2i and j). Because super-resolution imaging enables more precise visualization of size and structural features, the crosslinking-associated increase in fiber thickness can be directly observed in these images.

The averaged Raman spectra of collagen fibers obtained from the sclera and choroid of young and aged mice reveal differences in their relative density and structural characteristics. Although collagen fibers in young mice were distributed in closer proximity to the choroid, collagen fibers within the choroid itself were observed exclusively in aged mice (Fig. 2f, green dotted ellipsoid). Collagen fibers in the sclera of young mice exhibited the strongest Raman signal intensity, whereas those in the sclera of aged mice displayed a comparatively weaker signal while retaining a similar spectral profile. In contrast, collagen fibers in the aged choroid showed a distinct spectral shape compared to the scleral fibers. Specifically, the ratio of –CH_2_ to –CH_3_ stretching modes was higher in choroidal collagen than in scleral collagen (Fig. 2k). This finding suggests that choroidal collagen expressed in aged tissue may possess greater conformational flexibility than that in sclera, allowing increased vibrational activity of –CH_2_ groups in protein side chains.

### Increased lipid turnover in aged tissues

To quantify the lipid and protein metabolism in retinal tissues, animals were administered 20% D_2_O for 7 days and the turnover rates in retinal tissues were investigated using the DO-SRS and LC-MS approach.

In the DO-SRS analysis, both the lipid turnover rate and the detailed Raman spectra within the C-H stretching region were examined. Absolute quantification of Raman signals in the C-H stretching region was achieved by normalizing the target molecular signal against the water signal as an internal reference(38). The ratio of the C-D to the C-H vibrational signal serves as a measure of the anabolic activity of proteins or lipids.

The lipid turnover rate in both the outer segments of rod photoreceptor cells and the RPE layer was found to increase with age (Fig. 3a). Rod cell outer segments exhibited a comparatively lower baseline lipid turnover rate relative to other tissue compartments, and this inter-compartment difference was attenuated with aging. The mean lipid turnover rates in both the rod cell outer segments and the RPE layer were significantly elevated in aged retinas (Fig. 3b).

**Fig 3.**
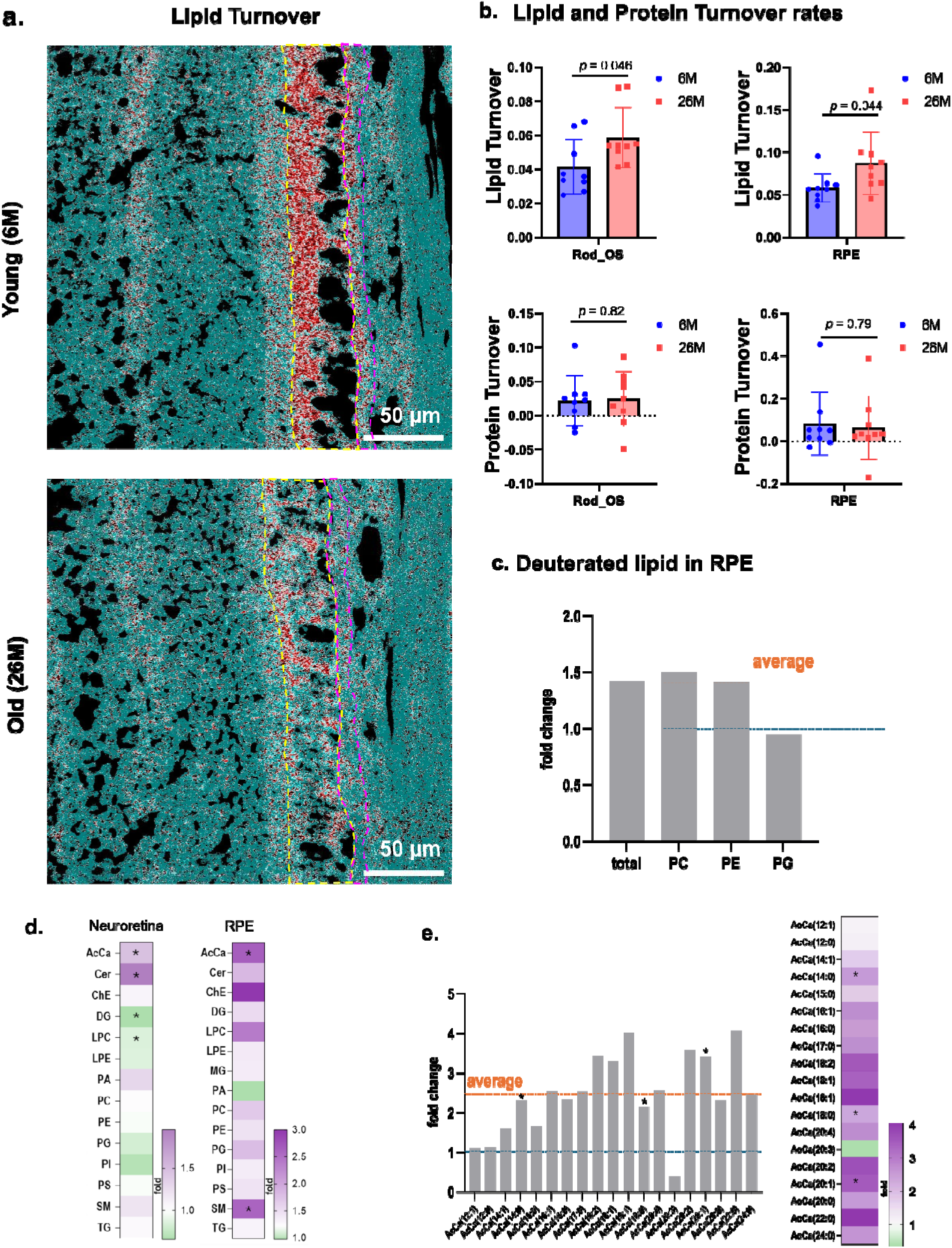
**a.** Newly synthesized lipid and old lipid distributions were measured using DO-SRS. In the outer segments of rod cells and RPE layer, lipid turnover rate was compared. **b.** Lipid turnover rate increase was detected in rod cell outer segment (yellow dashed line) and RPE layer (magenta dashed line) old retina sample. Protein turnover rate change did not show any significance. **c.** Consistent with the increased lipid turnover detected by DO-SRS, LC-MS analysis showed a 1.4-fold increase in deuterated lipids in the RPE, with PC and PE as the major contributors. **d.** In addition, AcCa increased with age in both tissues. In the RPE, AcCa levels were on average 2.5-fold higher in 26-month-old tissue than in 6-month-old tissue. **e.** Similarly, AcCa levels also increased in aged retinas, although to a lesser extent, reaching 1.4-fold. Taken together, these LC-MS data confirm higher lipid turnover in older tissues.

To further explore the increased lipid turnover in the RPE, we have performed LC-MS untargeted analysis on retinal samples isolated after D_2_O exposure. Our data show that in RPE there is a 1.4-fold increase of levels of deuterated lipids with predominant increase of PC and PE (Fig. 3c). Additionally, we have noted an increase of acyl carnitines (AcCa) in both tissues with age (Fig. 3d). Specifically, we detected an average 2.5-fold increase in the levels of AcCa in 26-month-old RPE when compared to 6-month-old tissues (Fig. 3e). Similarly, AcCa levels increased in aged retinas, although to a lesser extent (1.4-fold). Taken together, the LC-MS data support the imaging findings, indicating increased lipid turnover in the older tissues.

### Lipid compositional changes in rod cells and the RPE layer

HS-SRS image analysis combined with Penalized Reference Matching algorithm with SRS (PRM-SRS) yields detailed spectral information at the single-pixel level. By evaluating the spectral similarity between measured signals and reference spectra of known lipid subtypes, the relative abundance of each lipid species can be quantified. To ensure the accuracy of the similarity score analysis, water and protein vibrational contributions were subtracted from the hyperspectral images prior to analysis. The resulting lipid spectra across distinct tissue compartments were then systematically compared.

Specifically, aged retinal samples exhibited a significant reduction in the -CH_2_ stretching and - C=C- signals in the Raman spectra (Fig. S4). The most pronounced age-related spectral differences were observed in the rod outer segments. Following spectral unmixing of protein and lipid contributions, a significant decrease in the -CH_2_ and -C=C- signals of lipid was detected. (Fig. 4). Additionally, signals corresponding to the cholesterol ring and C=C stretching vibrations showed a tendency toward reduction in both tissue compartments.

**Fig. 4.**
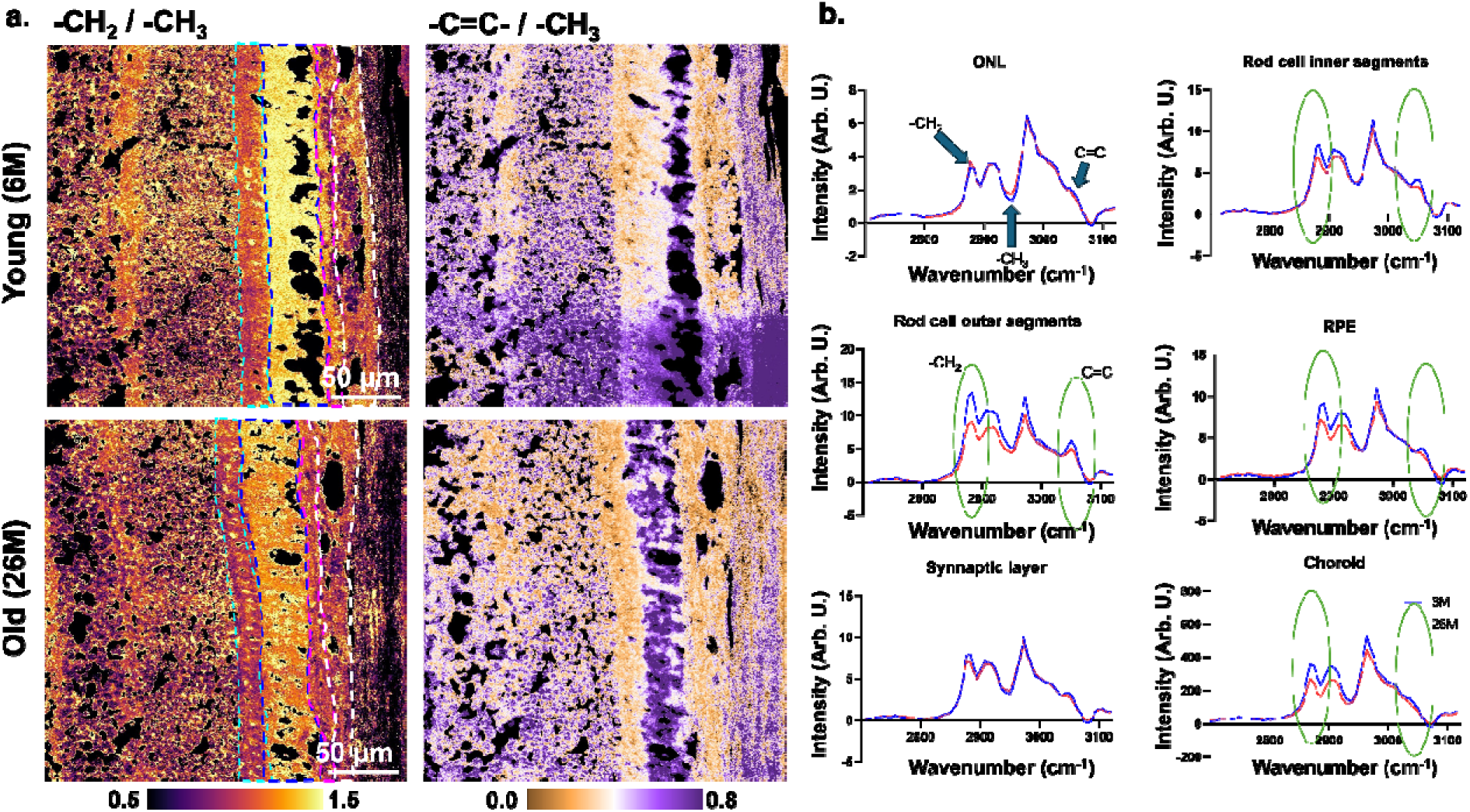
**a.** From HS-SRS image stacks, lipid chain length (−CH_₂_/−CH_₃_) and chain unsaturation (−C=C−/−CH_₃_) images were generated. Within the regions of interest (ROIs) corresponding to rod cell inner segments (cyan dotted line), rod cell outer segments (blue dotted line), the RPE layer (magenta dotted line), and the choroid (white dotted line), a decrease in both chain length and lipid unsaturation was observed in the retinas of aged mice. **b.** Averaged spectra from each ROI clearly illustrate the differences in lipid structural composition. Spectra from the ONL and synaptic layer showed no appreciable changes. The spectra from rod cell outer segments exhibited the most significant decreases in −CH_₂_ and −C=C− Raman signals. Rod cell inner segments, RPE, and choroid displayed a similar trend, albeit with lower statistical significance.

More subtle trends were detected in the rod inner and outer segments, RPE, and choroid, where decreases in both the -CH_₂_ vibrational signal and the C=C signal were evident but did not reach statistical significance. Despite these modest global changes, the most robust and quantifiable age-associated compositional alterations were identified in the rod inner segments and the RPE. In both regions, levels of triacylglycerols (TAGs) and cholesterol declined with age (Fig. 5a–d). In addition, sphingosine levels were significantly increased in the RPE of aged retinas (Fig. 5d).

**Fig. 5.**
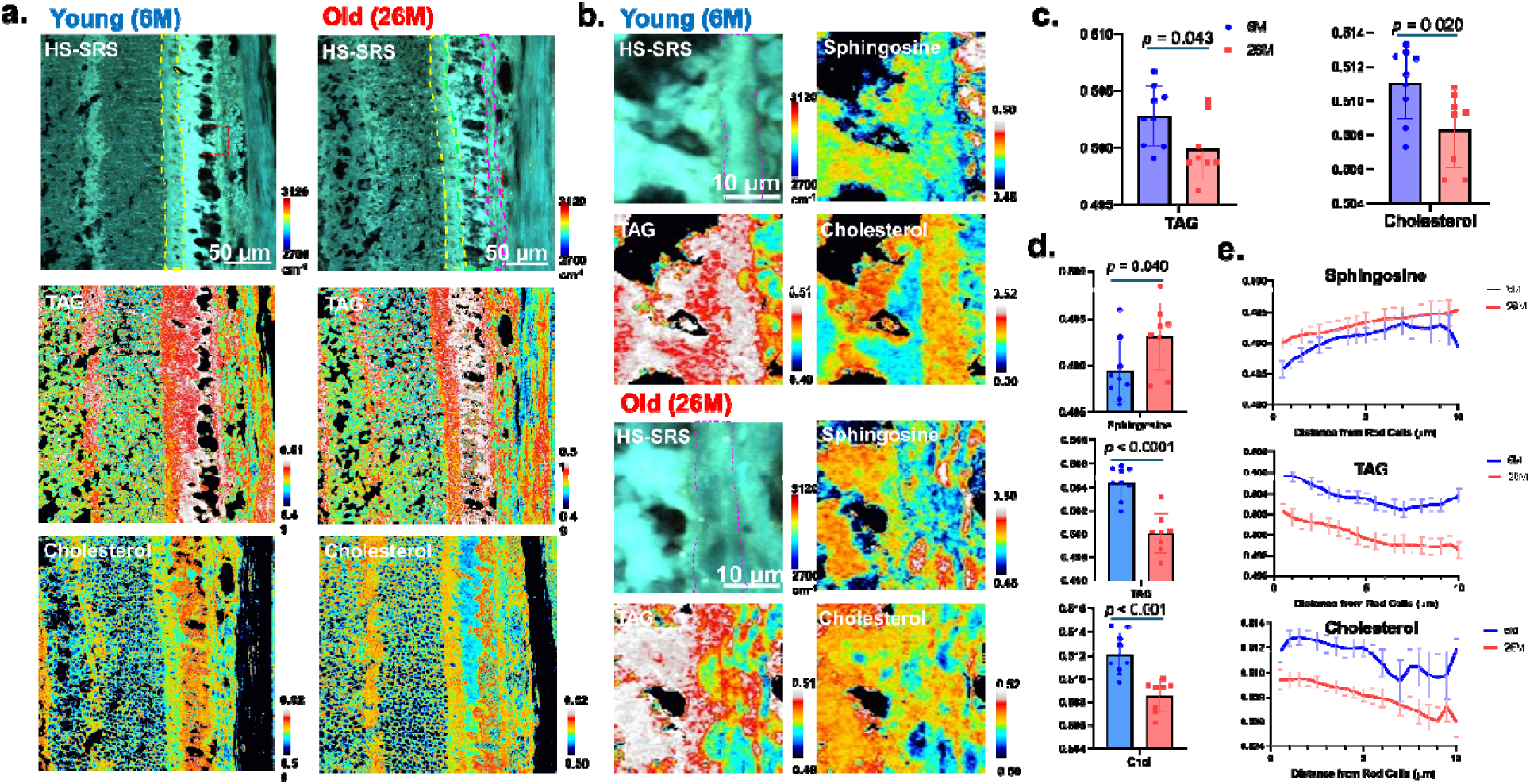
**a and b.** From hyperspectral SRS images, lipid compositions were analyzed using PRM-SRS method. TAG, and cholesterol distribution amount were detected. **c.** In rod cell inner segments (The region with yellow dashed line in HS-SRS), TAG and cholesterol decreased by aging. **d.** In RPE layers of two different aged retina samples, different levels of sphingosine, TAG, and cholesterol distribution amount were detected. The averaged lipid subtype contents show the sphingosine increase, TAG decrease and cholesterol decrease by aging in RPE layer. **e.** TAG and cholesterol did not show any aging-related spatial distribution changes. However, Sphingosine increase was mostly detected in the boundaries of RPE layer.

**Fig. 6.**
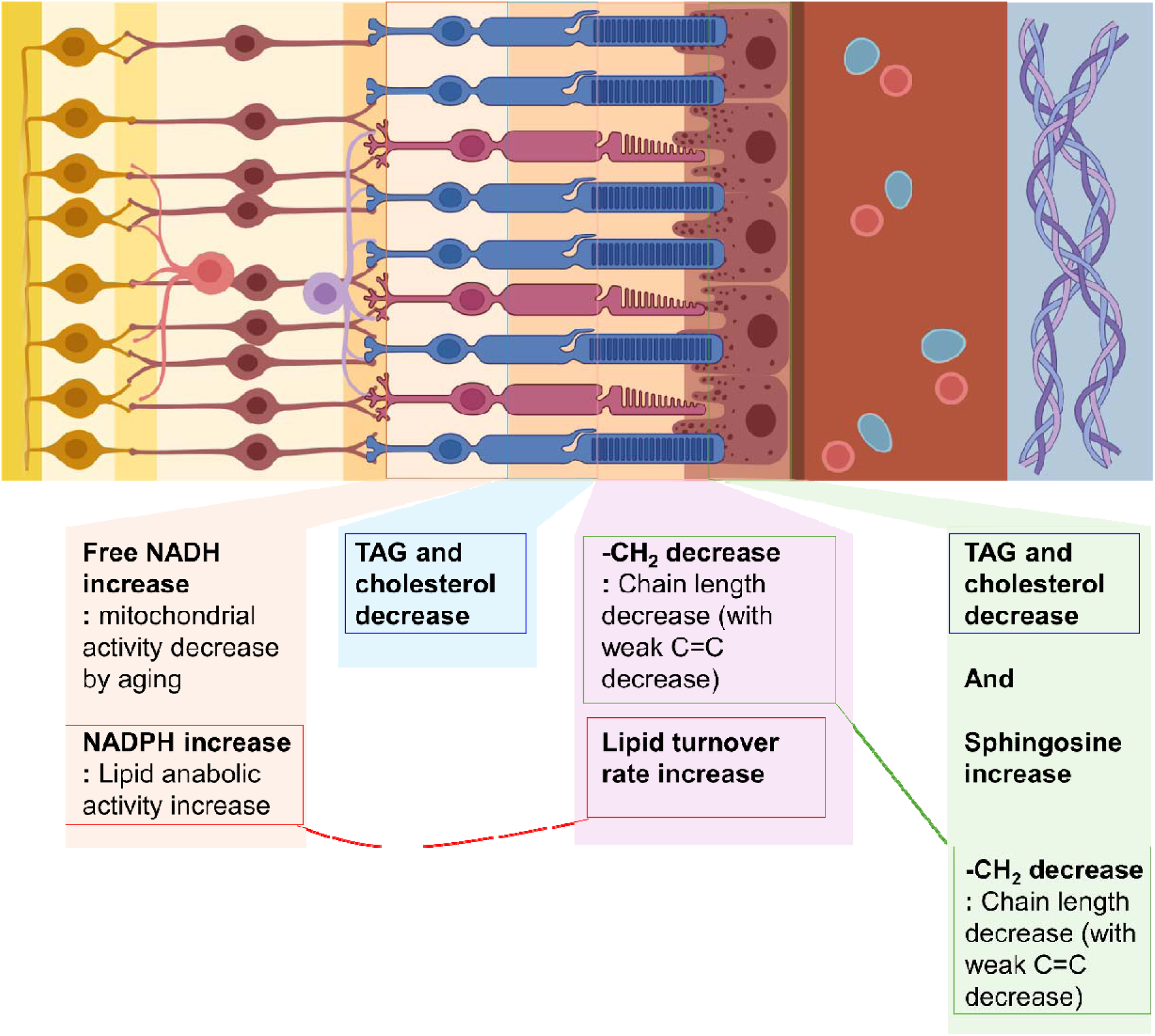
Summary of detected metabolic changes during aging. All identified metabolic signals are integrated and presented in a single schematic figure. The observed increase in NADPH levels is correlated with the elevated lipid turnover rate detected by DO-SRS. Consistent HS-SRS spectral differences were identified in both the RPE and rod cell outer segments. Furthermore, RPE and rod inner segments share comparable compositional alterations, characterized by a concurrent decrease in triacylglycerol (TAG) and cholesterol levels.

Using a spatial analysis approach analogous to that applied for collagen fiber characterization, the intra-layer distribution of lipid subtypes within the RPE was mapped (Fig. 5e). Both TAG and cholesterol exhibited a decreasing concentration gradient with increasing distance from the rod cells. In contrast, age-related changes in sphingosine levels were highly localized to the membrane-proximal region of the RPE layer. These spatially distinct distribution patterns likely reflect the specific biological functions of each lipid subtype within the retinal microenvironment.

Our untargeted lipidomic data analysis allowed us to corroborate and interpret some of the imaging data. First, we have noted higher levels of triglycerides in both retina and RPE (Fig. 5). Next, in both tissues, we found increased levels of ceramides and sphingomyelin in aging. Both types of lipids have a common core – sphingosine, therefore the data agrees with the imaging data. Interestingly, DG and LPC have inverted age-related levels change between the tissues (Fig. 3d). Specifically, both DG and LPCs are lower in neuroretina, while both are increased in aging RPE. These specific changes may reflect the metabolic needs of each tissue.

## DISCUSSION

The present study provides a comprehensive, spatially resolved characterization of age-associated metabolic, structural, and biochemical changes in the retina using multimodal nonlinear imaging. By combining SHG, FLIM, and hyperspectral SRS with isotopic labeling, the work moves beyond static descriptions to reveal aging as a dynamic process characterized by metabolic reprogramming, increased lipid turnover, and localized compositional remodeling.

A central observation is the shift in metabolic state within the ONL, as evidenced by fluorescence lifetime imaging of NADH, with the signal coming most likely from the photoreceptor mitochondria that are distributed quite evenly in the whole ONL(39). The increased relative abundance of free NADH and NADPH in aged retinal tissue suggests a decline in OXPHOS accompanied by a shift toward reductive and anabolic pathways. This interpretation aligns with prior studies demonstrating age-related mitochondrial dysfunction in both photoreceptors and RPE, including reduced respiratory capacity and increased oxidative stress(40–42). Given the high energetic demands of photoreceptors, which rely heavily on mitochondrial ATP production, such impairments are thought to play a central role in retinal aging and degeneration (41, 43–45). The observed increase in NADPH further suggests enhanced engagement of lipid biosynthesis and antioxidant systems, consistent with reports that aging cells reprogram metabolism to buffer oxidative damage while maintaining biosynthetic capacity (46, 47). We note that NADH and NADPH are spectrally indistinguishable and were resolved here on the basis of their distinct enzyme-bound fluorescence lifetimes(23); moreover, the free-to-bound NAD(P)H ratio is an indirect readout that can also be influenced by changes in the total NAD(H) pool size and in the availability of binding partners(66). These caveats are mitigated by our orthogonal DO-SRS measurements, which independently corroborate the elevated lipid anabolic activity inferred from the FLIM data.

In parallel with these metabolic changes, we observed a significant increase in lipid turnover rates in the rod photoreceptor outer segments and the RPE layer. While lipid accumulation has long been recognized as a hallmark of retinal aging, particularly in the context of lipofuscin formation and subretinal deposits (48, 49), our findings suggest that aging is not simply associated with passive lipid buildup but rather with dynamic remodeling. Indeed, the outer retina depends on continuous, lifelong lipid renewal: a substantial fraction of each photoreceptor outer segment is shed daily and phagocytosed by the RPE, and age-related dysregulation of this renewal cycle has been proposed as a central driver of outer retinal disease(51). This is in line with emerging perspectives in aging biology that emphasize dysregulated turnover rather than uniform metabolic decline(50–53). The increased lipid turnover detected here may reflect compensatory mechanisms aimed at maintaining membrane integrity in the face of cumulative oxidative damage, although such processes may ultimately become maladaptive. Consistent with this view, the accumulation of acylcarnitines detected in the aged RPE and retina is a recognized signature of mitochondrial overload and incomplete fatty acid β-oxidation(67, 68), suggesting that in the aged outer retina the delivery of lipid substrates outpaces mitochondrial oxidative capacity.

Consistent with this interpretation, hyperspectral SRS analysis revealed significant alterations in lipid composition with age, including reductions in triacylglycerol and cholesterol, alongside an increase in sphingosine within the RPE. These findings are broadly supported by previous lipidomic studies of the aging retina, which have documented disruptions in cholesterol homeostasis and enrichment of bioactive sphingolipids associated with cellular stress and apoptosis (51, 54–56). Sphingolipid metabolism, in particular, has been implicated in regulating photoreceptor survival and inflammatory signaling (57), suggesting that the observed increase in sphingosine may contribute to age-related vulnerability of retinal cells. Because sphingosine is generated through ceramidase-mediated hydrolysis of ceramide, the localized sphingosine accumulation imaged in the RPE, together with the elevated ceramides and sphingomyelins detected by LC-MS, is most consistent with broadly enhanced sphingolipid turnover in the aged outer retina. The pathological relevance of this axis is underscored by the finding that ceramide accumulation in AdipoR1-deficient mice drives photoreceptor loss, whereas its pharmacological inhibition improves photoreceptor survival and vision(69). Notably, the spatially resolved analysis performed here further demonstrates that these compositional changes are highly localized, with distinct gradients and microdomain-specific alterations within retinal layers. Although lipid accumulation in Bruch’s membrane and drusen is a hallmark of human retinal aging and AMD(54, 55, 63), such deposits are largely extracellular; the intracellular declines in TAG and cholesterol observed here, in the context of elevated turnover, may therefore reflect a shift of the aged RPE from lipid storage toward lipid flux and export. Furthermore, the reduced lipid acyl-chain length and unsaturation (-C=C-) observed in aged rod outer segments mirror the age-related decline in ELOVL2-mediated synthesis of docosahexaenoic acid and very-long-chain polyunsaturated fatty acids (VLC-PUFAs), which has been shown to functionally drive retinal aging(70). Notably, restoring VLC-PUFA content by intravitreal supplementation with the direct ELOVL2 product 24:5n-3 reverses aging-related visual decline in mice(71), suggesting that the compositional remodeling imaged here may be therapeutically actionable.

In addition to metabolic and biochemical remodeling, our results highlight pronounced structural changes in the sclera, characterized by increased collagen fiber density, anisotropy, and crosslinking in aged tissues. Moreover, increased levels of collagen were detected in the choroid, which may affect the function of this tissue. These findings are consistent with prior reports of age-dependent extracellular matrix remodeling and increased tissue stiffness, often attributed to processes such as non-enzymatic glycation and collagen crosslink formation(58, 59). Such biomechanical alterations may influence not only the structural integrity of the eye but also the diffusion of nutrients and metabolites across ocular tissues, thereby indirectly contributing to retinal metabolic stress.

Importantly, the integration of multimodal imaging with isotopic labeling in this study enables simultaneous assessment of molecular turnover, composition, and structure within intact tissue architecture. This represents a significant advance over conventional approaches that rely on bulk biochemical assays or single-modality imaging, which often lack spatial resolution or dynamic information. By resolving metabolic and compositional heterogeneity at the level of individual retinal layers, the present work provides a more nuanced understanding of how aging processes manifest within specific cellular microenvironments. Using LC-MS, we have shown that lipid turnover in the aged eye increases predominantly in the RPE, with a smaller increase in photoreceptors, suggesting a central role for the RPE in maintaining lipid homeostasis during aging.

These findings have direct implications for age-related retinal diseases, particularly age-related macular degeneration (AMD). AMD is characterized by dysfunction of the RPE, accumulation of lipid-rich deposits (drusen), mitochondrial impairment, and chronic oxidative stress(60–62). The metabolic shift away from OXPHOS, increased lipid turnover, and altered lipid composition observed in this study closely parallel key features of AMD pathogenesis. In particular, dysregulation of cholesterol and sphingolipid metabolism has been implicated in drusen formation and inflammatory activation in the RPE(55, 63). Moreover, the spatially localized increase in sphingosine within the RPE may reflect early biochemical changes that predispose these cells to degeneration. Together, these results support the notion that age-related metabolic reprogramming and lipid dysregulation are not only hallmarks of normal retinal aging but also potential drivers of disease progression in AMD.

In summary, this study establishes a multimodal, spatially resolved framework for interrogating retinal aging that integrates structural, metabolic, and chemical information within intact tissue. These methodological advances provide a powerful platform for dissecting microenvironment-specific changes that are not accessible through conventional bulk or single-modality approaches. At the same time, the interpretation of these findings should consider the limitations of ex vivo analysis, indirect metabolic readouts, and model-dependent spectral unmixing, as well as the use of a murine system without direct functional validation. Despite these constraints, the convergence of metabolic decline, lipid dysregulation, and structural remodeling observed here closely mirrors key features implicated in age-related retinal diseases, particularly AMD, underscoring the potential of this approach to inform future mechanistic studies and therapeutic strategies.

## Methods

### Multimodal Imaging Setup

The system is based on an SRS setup. A custom-built upright laser-scanning microscope (Olympus) equipped with a 25× water-immersion objective (XLPLN, WMP2, 1.05 NA, Olympus) was utilized for near-IR throughput. A synchronized pulsed pump beam (tunable 720–990 nm wavelength, 5–6 ps pulse width, 80 MHz repetition rate) and a Stokes beam (1032 nm wavelength, 6 ps pulse width, 80 MHz repetition rate) were supplied by a picoEmerald system (Applied Physics & Electronics) and coupled into the microscope. The pump and Stokes beams were collected in transmission by a high-NA oil condenser (1.4 NA). A high-O.D. shortpass filter (950 nm, Thorlabs) was used to completely block the Stokes beam and transmit only the pump beam onto a Si photodiode to detect the stimulated Raman loss signal. The output current from the photodiode was terminated, filtered, and demodulated by a lock-in amplifier at 20 MHz. The demodulated signal was fed into the FV3000 software module FV-OSR (Olympus) to reconstruct images during laser scanning. Retinal tissue images were acquired at a resolution of 1024 × 1024 pixels. The pixel dwell time was 20 μs. MPF, SHG, and FLIM were integrated into the SRS microscope to image the same region of interest using different modalities. Signals were collected using a photomultiplier tube (PMT), and FLIM measurements were performed by adding a TCSPC module to the computer for signal collection.

### FLIM Analysis

The FLIM data were processed using custom-built Python scripts for phasor plot(24) and lifetime curve fitting. As described in Fig. S1, a two-step analysis approach was employed. First, an uncalibrated phasor diagram was generated. From this phasor diagram, pixels representing different chemical environments were clustered using the Mean Shift algorithm(64). Following clustering, the photon-counting histogram of each pixel cluster was averaged and fitted with a biexponential decay model. By using these fitting results as initial values for the precise fitting of individual pixel lifetime curves, a detailed two-component analysis of free and enzyme-bound flavin and NADH was achieved. Finally, the lifetime of enzyme-bound NADH was used to calculate the NADPH/NADH ratio(23).

### SHG Collagen Analysis

Collagen in SHG images was identified by applying an intensity-based segmentation approach, and collagen density was calculated as the proportion of collagen-positive pixels within each ROI. Fiber thickness and orientation were then assessed using Otsu-based thresholding in combination with orientation tensor analysis. The alignment metric shown in the supplementary figures describes the spread of local fiber orientation angles, where lower values correspond to a more uniform and parallel collagen fiber arrangement. SHG signal intensity was evaluated only across samples imaged with the same acquisition parameters, and no adjustments were made for effects related to polarization sensitivity or imaging depth. Thus, throughout this study, “collagen density” denotes collagen area fraction rather than an absolute measure of collagen concentration.

### Image analysis for DO-SRS and PRM-SRS

Lipid subtype composition was determined using the PRM-SRS method(65). Hyperspectral images covering the C-H vibration region (2700-3120 cm^-1^) were acquired at 6 cm^-1^ intervals, yielding a stack of 71 frames per field of view. Spectral similarities between the acquired data and a reference library, identical to that used in the previous PRM-SRS study, were calculated to identify the lipid subtypes present in each sample. The results were expressed as normalized ratiometric images for each lipid subtype, computed as [target lipid] / ([target lipid] + [PE]).

Hyperspectral images in the C-D vibration region (2000–2300 cm^-1^) were acquired at 6 cm^-1^ intervals, producing 51 frames per stack. From these stacks, newly synthesized lipid images (2135 cm^-1^) and background images (2000 cm^-1^) were extracted. To correct baseline intensity offsets, the background images were subtracted from the newly synthesized lipid images. The corrected images were then divided by the existing lipid channel image, derived from the C-H hyperspectral stack, to convert absolute new-lipid signals into lipid turnover rate maps.

### Mouse Retina Tissue Slice Preparation

All animal procedures were conducted with the approval of the Institutional Animal Care Committee (IACUC) at the University of California, Irvine, under AUP #23-063. Male albino mice, aged 6 and 26 months, were acquired from The Jackson Laboratory. The mice were housed in the vivarium at the University of California, Irvine, under a standard 12-hour light (<150 lux)/12-hour dark cycle. They were provided with a standard soy protein-free rodent chow diet (Envigo Teklad 2020X) *ad libitum*. Tissues were collected between 10AM and 12PM to assure similar conditions in all animals.

For D_2_O probing experiments, mice that consumed 20% D_2_O for 7 days were anesthetized using isoflurane and subsequently euthanized. Following euthanasia, mouse eyes were enucleated and immersed in 4% paraformaldehyde (PFA) dissolved in phosphate-buffered saline (PBS, pH 7.4) for 2 hours at 4°C. The cornea, lens, and vitreous were then meticulously removed, and the remaining eyecups were fixed in 4% PFA overnight at 4°C. The cryoprotected eyecups were then embedded in Tissue-Tek OCT (Sakura, Torrance, CA) and rapidly frozen on a conductive metal block cooled by dry ice. Cryosectioning was performed using a cryostat to obtain 12 µm-thick serial sections of the retina.

### Lipidomic analysis

#### Lipid extraction

Lipid extractions were performed according to the methodology of Bligh and Dyer (PMID: 13671378). In brief, the tissue was homogenized in 200 μL water, transferred to a glass vial, and 750 μL 1:2 (v/v) CHCl3: MeOH was added and vortexed. Then 250 μL CHCl3 was added and vortexed. Finally, 250 μL ddH2O was added and vortexed. The samples were centrifuged at 3000 RPM for 5 min at 4 °C. The lower phase was transferred to a new glass vial and dried under nitrogen stored at -20 °C until subsequent lipid analysis.

#### LC-MS/MS

Separation of lipids was performed on an Accucore C30 column (2.6 μm, 2.1 mm × 150 mm, Thermo Scientific). The Q Exactive MS was operated in a full MS scan mode (resolution 70,000 at m/z 200) followed by ddMS2 (17,500 resolution) in both positive and negative mode. The AGC target value was set at 1E6 and 1E5 for the MS and MS/MS scans, respectively. The maximum injection time was 200 ms for MS and 50 ms for MS/MS. HCD was performed with a stepped collision energy of 30 ± 10% for negative and 25%, 30% for positive ion mode with an isolation window of 1.5 Da.

## Data analysis and post-processing

Data were analyzed with LipidSearch 4.2.21 software. Only peaks with molecular identification grade: A or B were accepted (A: lipid class and fatty acid completely identified or B: lipid class and some fatty acid identified). The relative abundance of each lipid species was obtained by normalization to the total lipids intensity.

## Acknowledgement

We thank Drs. C. Metallo, P. Adams for meaningful discussions.

## Funding

This work was in part supported by the United States National Institutes of Health (NIH) grants (R01AG086548, R01GM149976, U01AI167892, R01HL170107, R01NS111039, R21NS125395, U54DK134301, U54 HL165443), UCSD Startup funds, Sloan Research Fellow Award, and CZI DAF2023-328667 Award (all to L.S.). D.S.-K. was supported by the NIH National Eye Institute U01 (EY034594), P30 (EY034070) and by the RPB / Dr. H. James and Carole Free Catalyst Award for Innovative Research Approaches for AMD. The Gavin Hebert Eye Institute at University of California, Irvine was provided by an unrestricted grant from Research to Prevent Blindness.

## Author contributions

Conceptualization: L.S.& D.SK. Methodology: H.J. Resources: S.W., F.G. Investigation: L.S.,D.SK., H.J. Visualization: H.J. S.W., F.G. Supervision: L.S., D.SK Writing—original draft: H.J. & L.S. Writing—review and editing: L.S. D.SK , H.J., S.W., F.G.

## Competing interests

The authors declare no competing interests.

## Data, code, and materials availability

All imaging data, raw or processed ones are properly stored at UC San Diego and UC Irvine campus computers and external storage disks and are available upon request. All code is available for downloading from Github: https://github.com/lingyanshi2020

## Notes

### Competing Interest Statement

The authors have declared no competing interest.

