## Supplemental Figures for "Multimodal optical imaging reveals spatial metabolic heterogeneity in the aging retina"

**Supplementary Figures**

**Spatially Resolved Metabolic Alterations in the Aging Retina Revealed by Nonlinear Multimodal Imaging**

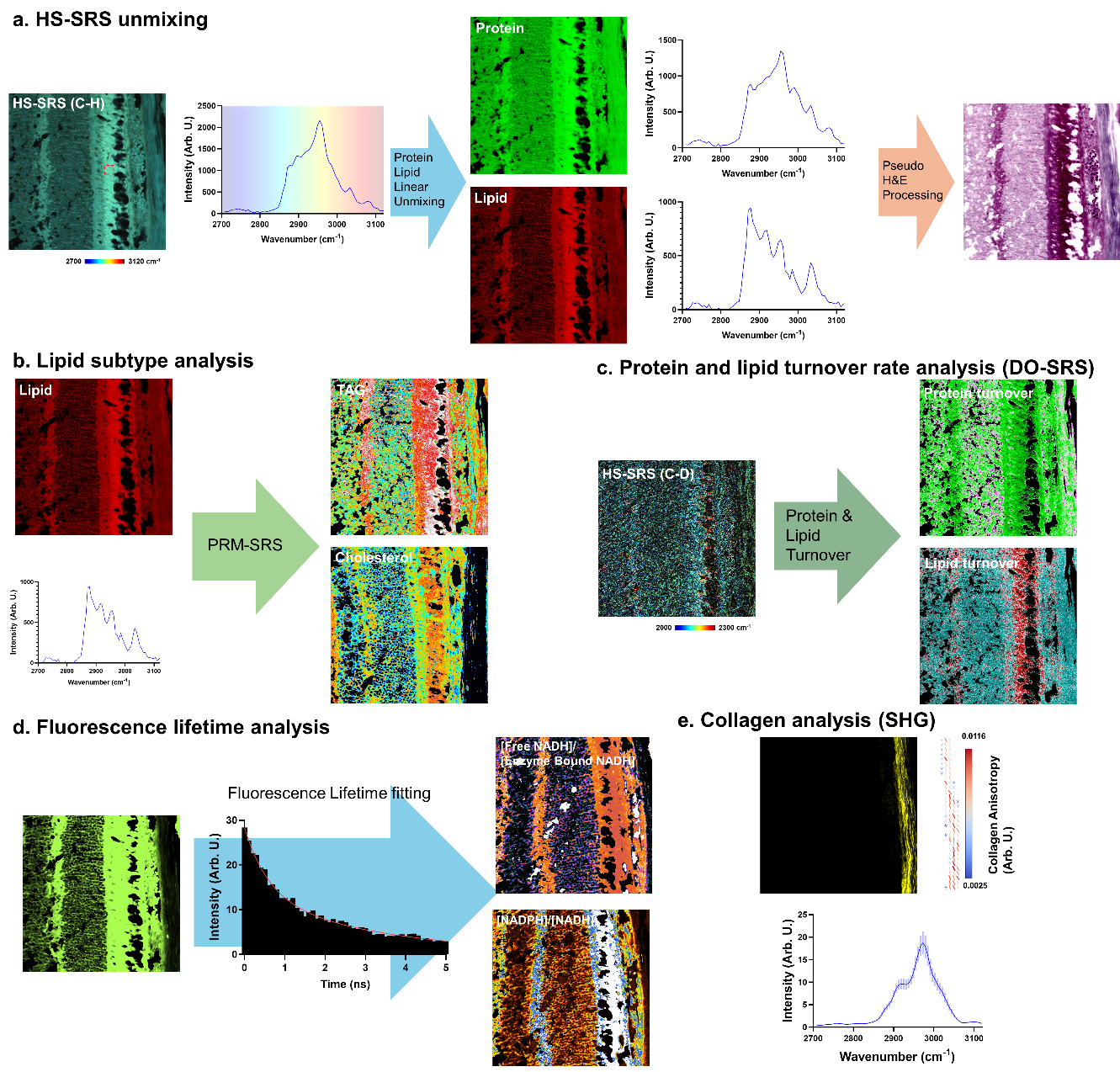

**Fig S1. Image analysis pipeline. a.** HS-SRS image stacks were linearly unmixed to obtain separate protein and lipid signal images. Pseudo-H&E images were then generated based on the intensity ratio of protein to lipid. **b.** From the unmixed lipid-channel HS-SRS images, quantitative spatial distributions of TAG and cholesterol were calculated using PRM-SRS. **c.** From the C–D channel HS-SRS images, newly synthesized lipid and protein images were extracted. Dividing these images by the corresponding pre-existing lipid and protein images yielded protein and lipid turnover rate maps. **d.** FLIM analysis was performed to measure free NADH and NADPH levels. **e.** From the SHG images, collagen distribution was mapped. Raman spectral signals averaged over ROIs within collagen fibers enabled structural analysis of collagen.

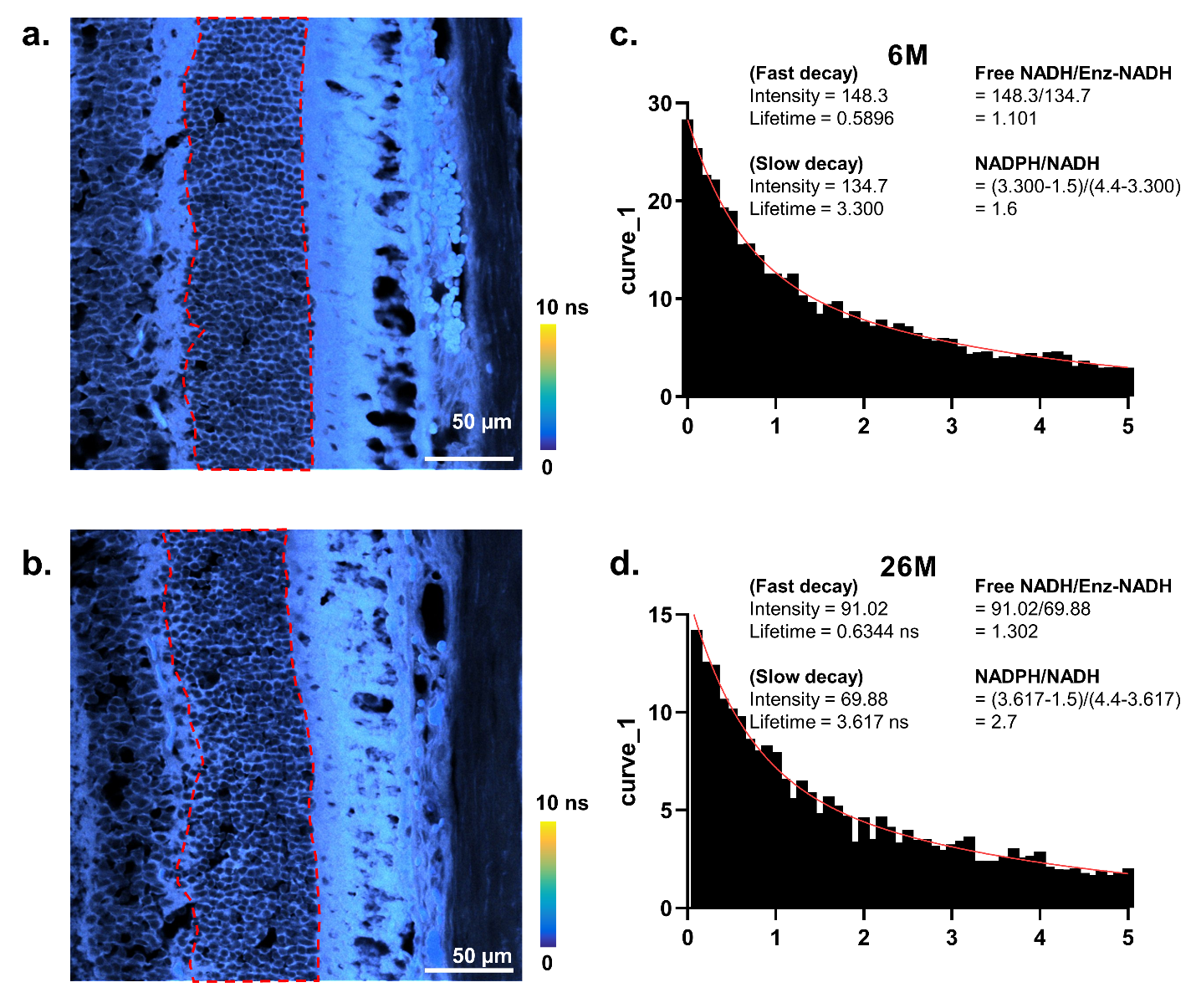

**Fig S2. FLIM data analysis in the ONL. a and b.** FLIM signals were extracted from measured FLIM images within each region of interest (ROI) in the retina and subsequently averaged. Signals originating specifically from the outer nuclear layer (ONL, red dotted region) were selectively averaged following bi-exponential decay fitting. **c and d.** From the fitting parameters obtained using the bi-exponential decay model, components corresponding to the fast and slow exponential decay curves were separately determined. Based on these fitted parameters, the relative levels of free NADH, enzyme-bound NADH, and NADPH were calculated as described.

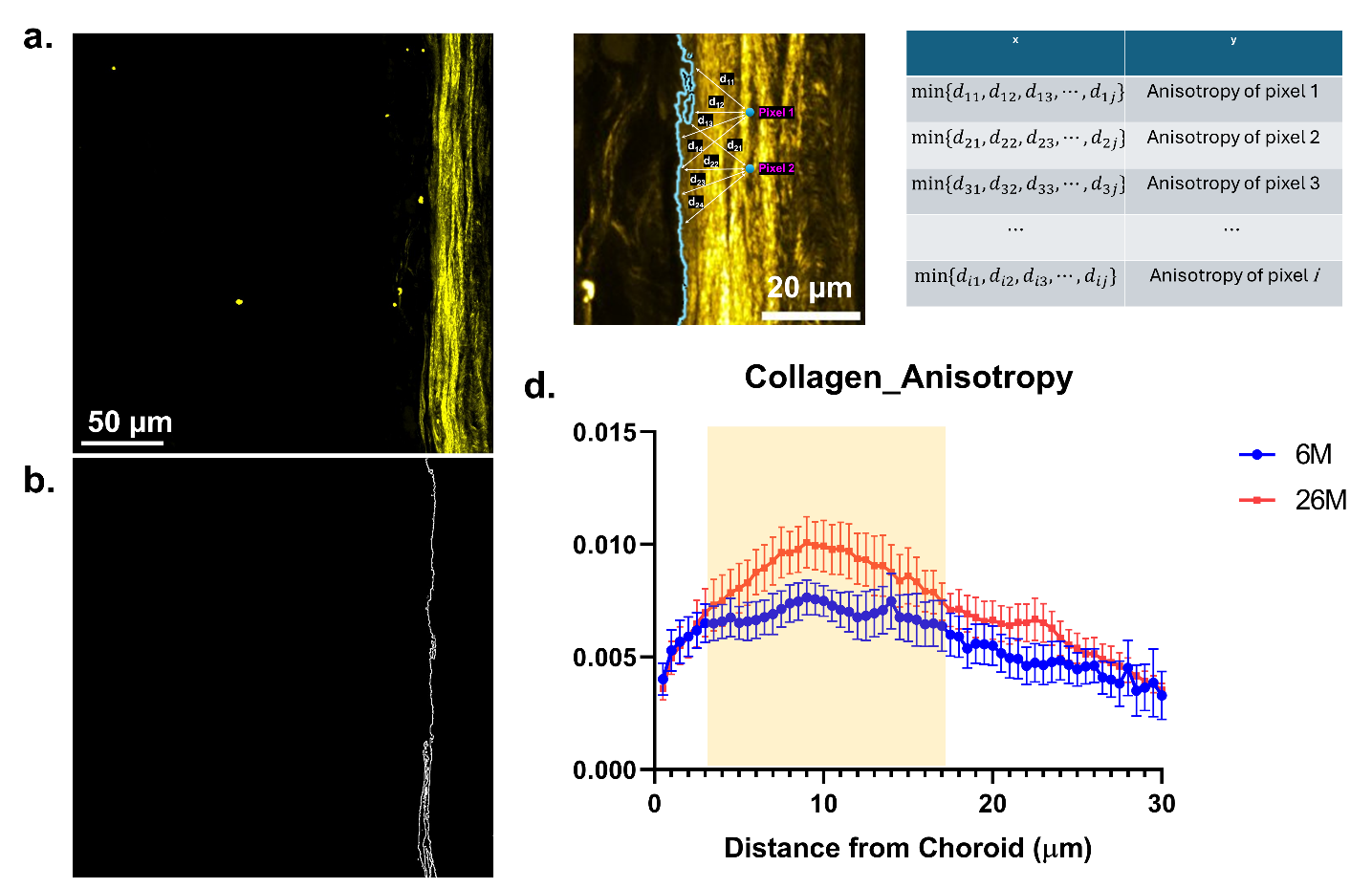

**Fig S3. Distance-dependent analysis of collagen fiber anisotropy. a.** Representative SHG image of an aged retinal sample. **b.** The boundary between the sclera and choroid was defined using an edge detection algorithm. **c.** For each pixel corresponding to a collagen fiber in the SHG image, the distance to every pixel along the defined boundary was computed, and the minimum distance was recorded along with the corresponding anisotropy value of that pixel. **d.** The resulting dataset of minimum distances and anisotropy values was compiled and represented as a single scatter plot.

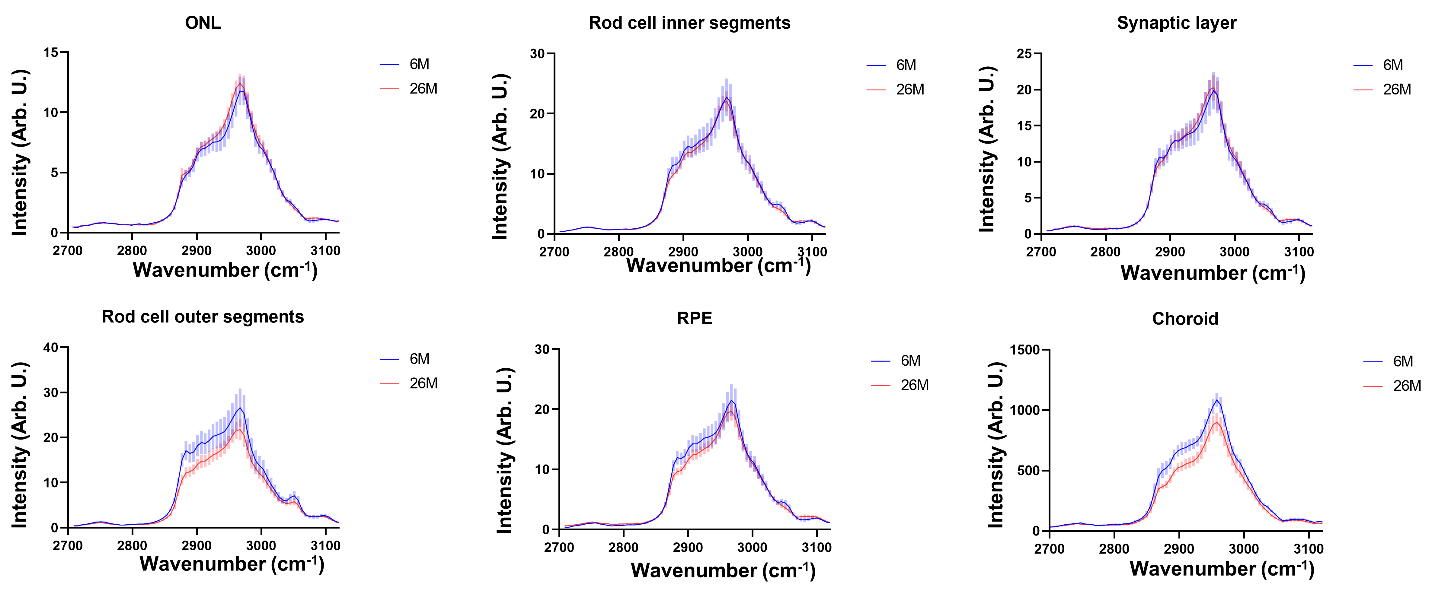

**Fig S4. C-H stretching Raman spectra analysis.** Raw C–H stretching spectra across retinal layers, in which meaningful spectral differences were observed exclusively in the rod outer segments, RPE, and choroid. Rod cell inner segments had the tendency of the similar signal decrease with very low significance.
